# Counting giraffes: A comparison of abundance estimators on the Ongava Game Reserve, Namibia

**DOI:** 10.1101/2021.09.28.459593

**Authors:** Bonenfant Christophe, Stratford Ken, Périquet Stéphanie

**Author notes:** (Corresponding author) Université de Lyon, F-69 000, Lyon; Université Lyon 1; CNRS, UMR 5558, Laboratoire de Biométrie et Biologie Évolutive, F-69 622, Villeurbanne, France.

## Abstract

Camera-traps are a versatile and widely adopted tool for collecting biological data for wildlife conservation and management. While estimating population abundance from camera-trap data is the primarily goal of many projects, the question of which population estimator is suitable for analysing these data needs to be investigated. We took advantage of a 21 day camera-trap monitoring period of giraffes (*Giraffa camelopardalis angolensis*) on the Ongava Game Reserve (Namibia) to compare capture-recapture (CR), rarefaction curves and *N*-mixture estimators of population abundance. A marked variation in detection probability of giraffes was observed both in time and between individuals. Giraffes were also less likely to be detected after they were seen at a waterhole (mean daily visit frequency of *f* = 0.25). We estimated the population size to be 119 giraffes (*C*_*v*_ = 0.10) using the most robust reference estimator (CR). All other estimators deviated from the CR population size by *ca*. −20 to > +80%. This was due the fact that these models did not account for the temporal and individual variations in detection probability. We found that modelling choice was much less forgiving for *N*-mixture models than CR estimators because it leads to very variable and inconsistent estimations of abundance. Double counts were problematic for *N*-mixture models, challenging the use of raw counts (*i*.*e*. when individuals are not identified) at waterholes, to monitor the abundance of giraffe or of other species without idiosyncratic coat patterns.

## Introduction

The on-going development and large-scale deployment of camera trapping technology offers a promising and appealing way for ecologists to collect a variety of biological data at an unprecedented scale and speed (Swanson et al. 2015). Habitat use, activity patterns and population abundance are now frequently studied using camera trap data (O’Connell et al. 2011; Trolliet et al. 2014). Sampling a population with camera-traps is indeed particularly useful and efficient (Wearn & Glover-Kapfer 2019), even more so for species with idiosyncratic coat patterns from which individual identification is possible (*e*.*g*. Jackson et al. (2006); Karanth & Nichols (1998); Stratford & Stratford (2011)). Camera trap data are increasingly used to estimate population abundance (Burton et al. 2015; Gilbert et al. 2021) but such data come with specific problems. Detection rate is not perfect, and sampling design and effort are likely different from physical captures (Hamel et al. 2013; Gilbert et al. 2021). While obtaining unbiased estimates of abundance is of central importance for conservation and wildlife management to set appropriate goals and policies (Anderson 2001), the suitability of the currently available population abundance estimators for camera-trap data remains to be evaluated empirically.

The underestimation of population abundance has been reported in the wild for a long time (Strandgaard 1967; Apollonio et al. 2010). This phenomenon arises because an unknown proportion of animals are missed during surveys, *i*.*e*. animal detection is not perfect. Imperfect detection is the main reason why detection probability of individuals underpins most population abundance estimators (Seber 1982; Schwarz & Seber 1999). Past empirical studies showed how variable detection probability could be in both time and space (Otis et al. 1978). For instance, detection probability was reported to increase with habitat openness (Choquenot 1995), vary between con-specifics with different behavioural repertoires (*i*.*e*. personalities, see Le Cœur et al. 2015, for an example on Siberian chipmunk *Tamias sibiricus*), decrease with the distance of animals from the observer (Burnham et al. 1980; Buckland et al. 2000), between observers themselves depending on their experience or motivation in spotting animals (Collier et al. 2007), and between camera trap brands or orientation (Rovero et al. 2013).

Accounting for these intrinsic and extrinsic sources of detection heterogeneity has profound consequences for the accuracy and precision of population abundance (Veech et al. 2016). Currently, only a handful of population abundance estimators can account for the multiple sources of variability in detection probability, and most derive from either distance sampling (DS) and capture-recapture (CR). Both families of estimators can accommodate detection rate for known sources of variability like time of the year, habitat type, or sex and age of individuals (Pollock 1980; Schwarz & Seber 1999). However, only the CR approach can model unmeasured or unknown sources of heterogeneity. The reason why these two methods are not systematically implemented in the field is due to serious practical limitations. CR requires a substantial proportion of the population to be recognizable: for instance Strandgaard (1972) recommended that up to 2/3 of a roe deer population should be marked to obtain meaningful results. In addition, the capture and marking of wild animals can raise ethical questions for endangered species. DS on the other hand, is quite sensitive to the sampling designs (*e*.*g*. linear transects and coverage), and is sometimes difficult to carry out in dense tropical forests of Africa (Duckworth 1998), or when human disturbance induces behavioural responses (see Elenga et al. 2020, on duikers). In other words, these two reference methods for estimating animal abundance can rapidly become prohibitively expensive, time consuming and difficult to implement at large spatial scale for wildlife managers (Morellet et al. 2007).

By seeking to keep implementation costs low, practitioners often make use of easier-to-implement, cheaper methods to monitor wildlife populations at spatial scales compatible with wildlife management (Morellet et al. 2007). This choice often comes at the costs of using estimators with less flexibility in accounting for variability in detection rate. For instance, the catch-per-unit effort (Leslie & Davis 1939) or the rarefaction curves (Petit & Valiére 2006) can return an estimate of population size from unmarked animals, but both assume constant detection rates for all individuals over the sampling period. A noticeable exception is the *N*-mixture model (Royle 2004), which allows the separation of population size from detection probability using repeated counts of animals in time and space. The robustness and accuracy of *N*-mixture abundance estimators is, however, frequently questioned (Kéry 2018).

For decades in large African national parks, a common practice has been to monitor wildlife using indices of population abundance of large herbivore species from direct (observation of animals) or indirect observations (observation of signs like tracks, faeces) (Jachmann 2002; 2012). Such indices can be obtained through road transects counts (with visibility issues), aerial counts (with visibility issues and costs), and waterhole counts of various duration (with the risk of missing water independent species). The underlying assumption of a constant detection rate has been advanced to be the main reason for indices of population abundance to fail at monitoring wildlife abundance reliably (Anderson 2001). However, these indices might be suitable for use by managers following a validation test against a reference method (Morellet et al. 2007). While several studies show that not accounting for detection variability can indeed bias population abundance estimates (Dail & Madsen 2011), the magnitude and direction of this bias is seldom quantified empirically.

The giraffe (*Giraffa camelopardalis ssp*.) is a charismatic species of conservation significance with decreasing populations in many parts of Africa (O’connor et al. 2019). The assessment of local populations’ conservation status and their long-term viability are however hampered by the many different ways abundance has been estimated between study areas. Here, we propose to take advantage of waterhole monitoring with camera traps on the Ongava Game Reserve, Namibia, to compare six population size estimators to characterize the biases associated with spatial, temporal and individual variability in detection rates. Being water dependent but with a capacity to spend several days without drinking, individual giraffes typically come to drink every two or three days (Shorrocks 2016). This behaviour can potentially generate variation in detection probability once individuals has visited a waterhole, *i*.*e*. an individual seen on a given day will be less likely to be seen on the following day. It is also known that males and females have different behaviours and resource requirements (see Gaillard et al. 2003, for examples in different large herbivore species), therefore the frequency of waterhole visit might differ between sexes (Shorrocks 2016).

A practical advantage of using giraffe as a study species is that one can use its idiosyncratic patterns to uniquely identify individuals from photographs, and then apply CR estimators to evaluate population abundance (Brown et al. 2019). This offers a unique opportunity to quantify the impact of detection heterogeneity on population size estimates, and to assess the relevance of simpler indices of abundance to monitor giraffe (and other species) populations. We compared the abundance estimates obtained from tried and tested CR methodologies, with *N*-mixture estimates, rarefaction curves, and raw count data (by observers) on the Ongava Game Reserve in 2016.

## Material and Methods

### Study area

Ongava Game Reserve (OGR) is located in Namibia, covering an area of approximately 300km^2^ immediately to the south of Etosha National Park with a common boundary on Ongava’s north side (Fig. 1). OGR is enclosed by electrified fences preventing movement of ungulates in and out the reserve. The habitat is termed *Karstveld*, with vegetation primarily (*Colophospermum mopane*) shrub and woodland, with some areas savannah-like. OGR’s relief is mostly dolomite hills, with a few small open plain areas and a well-defined ridge and small mountains in the central and northern part of the reserve. The weather zone for the reserve is typical for semi-arid northern Namibia, with an average annual rainfall of 380mm (see Stratford & Stratford 2011, for further details). There are several natural dams on the reserve, although most of these only contain water during the rainy season (January - April). During the dry season (May to December) water is only available at 12 artificial waterholes.

**Fig. 1.**
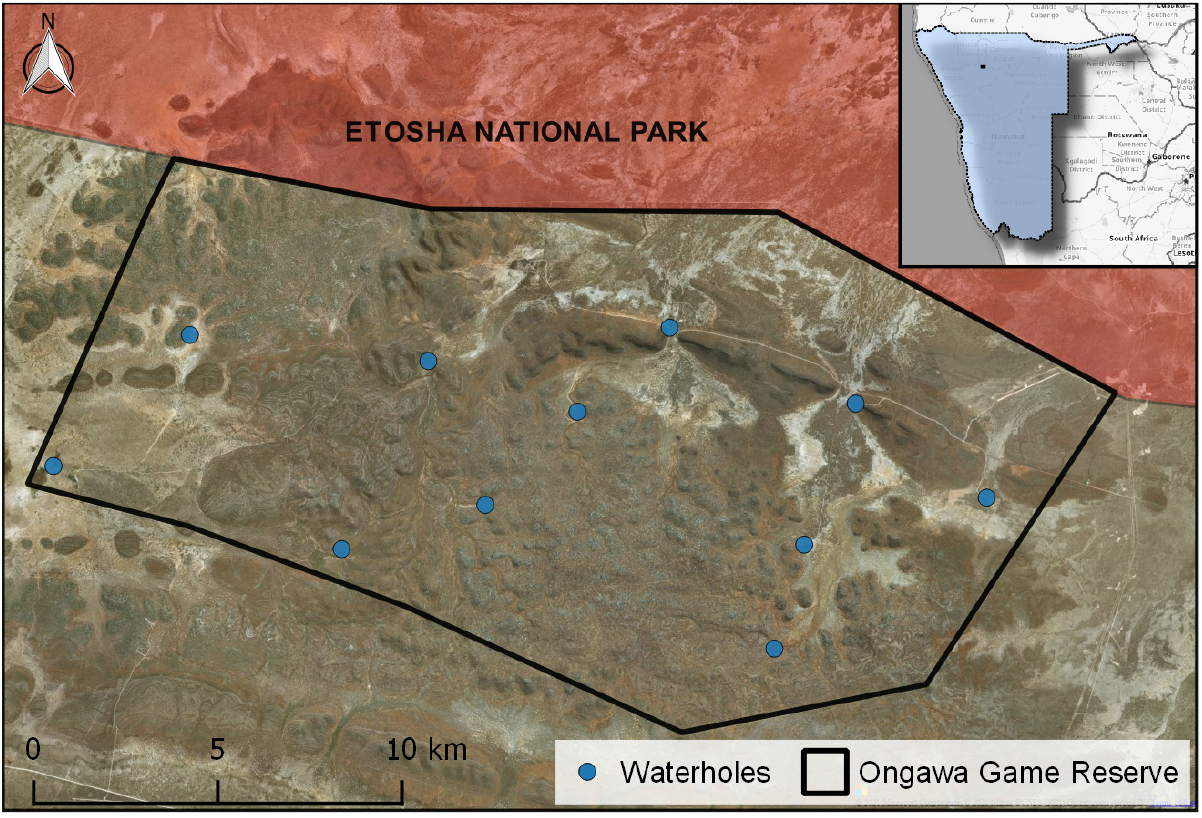
Spatial distribution of waterholes surveyed in 2016 with camera traps to monitor wildlife abundance at Ongava Game Reserve, Namibia. We extracted abundance data for giraffe (*Giraffa camelopardalis angolensis*) to be apply to different estimators of giraffe population size.

### Count data

From the 8^th^ to the 27^th^ of September 2016 (a total of 20 days), between three and eight camera traps (®Reconyx RC-55 and HC-500 and ®Bushnell Trophy series) were deployed at each waterhole to monitor their usage by wildlife (see Table S1). Each camera was mounted inside a stainless-steel protection case bolted to a tree or a pole within 10–15m of the waterhole. Reconyx cameras were set to record a sequence of 10 images with a delay of 30 seconds between sequences, while Bushnell cameras recorded sequences of 3 images with a delay of 15 seconds. We extracted all images containing giraffe and their associated metadata (date and time).

The camera traps yielded a total of 30 913 giraffe images. From these, 85 were discarded because the date and time of capture recorded by the camera were wrong. When possible, individual giraffe were manually identified in each image based on their unique coat patterns with the help of HotSpotter software (Crall et al. 2013). Whenever a giraffe could not be identified from its coat patterns or with the help of other images in the sequence, it was labelled as unknown. Where possible, we recorded the age-class (adults, sub-adults and juveniles) and sex of each individual was recorded.

### Population size estimations

#### Capture-Recapture models

We built daily capture histories for each individual giraffe over the *t* = 20 days of the camera trap survey. We then analysed these capture histories with CR methods (Lebreton et al. 1992) in a Bayesian framework (see Kéry & Schaub 2011). Each giraffe observation at a waterhole is the product of survival (*ϕ*) and detection (*p*) probabilities, conditional on first observation. We modelled detection probability *p* on the logit scale as a function of time (i.e day, categorical variable with 19 levels), whether the individual has been seen at any waterhole the previous day or not (categorical variable with 2 levels), and of the total number of functioning cameras (covariate). We also included random effects of the individual 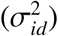 and of time 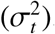. Because we could not identify the sex of two individuals, we treated sex as a latent Bernouilli variable *S* _*i*_ of parameter *π* corresponding to the population sex-ratio. We then entered *S* _*i*_ as an explanatory variable (categorical variable with 2 levels) of *p*.

Taken together, our set of fitted models covered the standard estimators for population size namely M_*t*_ (time effect), M_*th*_ (time and individual heterogeneity effects) and M_*tbh*_ (time, individual heterogeneity and behavioural effects: see Otis et al. 1978). We selected the statistically significant variables from the posterior parameter distributions and only kept variables for which 0 was excluded from the 95% credible interval. In our case, survival rate *ϕ* was of no biological relevance and kept constant in time and across individuals. We derived population sizes at each time step (*N*_*t*_) by dividing the number of giraffes detected at time *t* by the average detection probability *p* at time *t* + 1 (Burnham et al. 1987).

We considered the 20 *N*_*t*_ values as the realisation of an unobserved random variable of mean (*N*_*c*_), the estimated population size of giraffe, and of variance 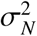, known as a Kalman filter (Kalman 1960; see Wang 2007 for an application in population ecology). We used an open population model because, despite OGR being thoroughly fenced, we could not completely ensure that all animals were always at risk of observation for the 20 days of survey. Several giraffe have been seen only once (*n* = 11), suggesting some individuals virtually leave the monitored population, becoming not longer at risk of observation.

#### Rarefaction curves

We also estimated population size using the rarefaction curves method (see Petit & Valiére 2006). Rarefaction curves have been used similarly for decades to estimate species diversity (Colwell & Coddington 1994). Over the course of the survey, the cumulative number of different giraffe seen at waterholes (hereafter noted *C*_*t*_) increased from day 1 to day 20 (see Fig. 2). Two different non-linear functions have been proposed in the literature for the case of population size estimation:

**Fig. 2.**
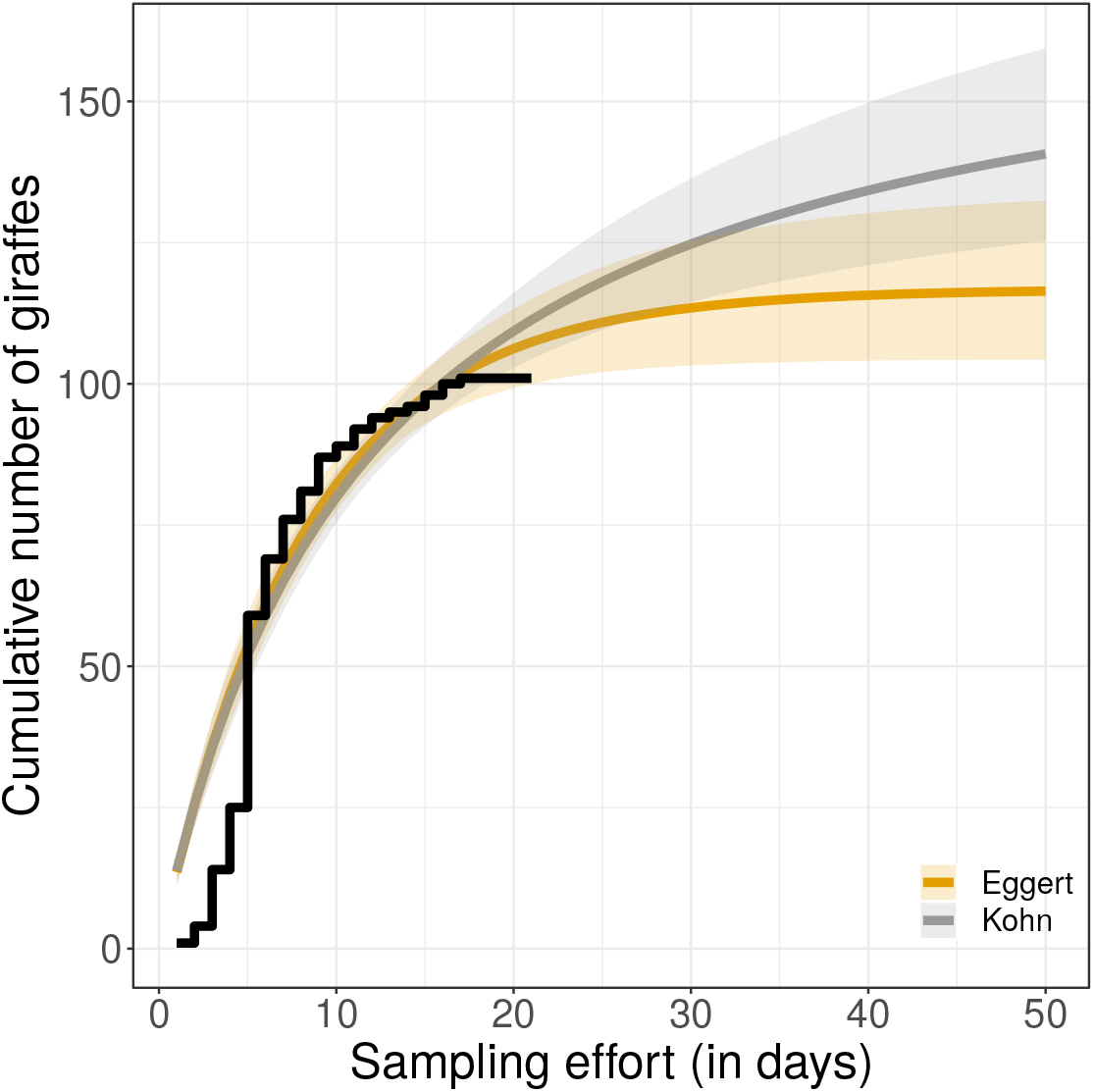
Rarefaction curves for individual giraffe (*Giraffa camelopardalis angolensis*) detected during the 21-day study period on Ongava Game Reserve, Namibia (step curve in black). Continuous lines and associated shaded areas represent predictions and credible intervals of rarefaction models. We fitted two rarefaction equations proposed by Eggert et al. (2003) and Kohn et al. (1999) to the problem of population size estimation using a Bayesian framework.

1. the hyperbolic function (Kohn et al. 1999): *C*_*t*_ = (*N*_*s*_ × *t*)/(*b* − *t*);
2. the exponential function (Eggert et al. 2003): *C*_*t*_ = *N*_*s*_ × (1 − *e*^−*c*×*t*^);

where *t* is time in days ranging from 1 to 20, and *b* and *c* are breakage parameters, *i*.*e*. the rate of decrease of the number of new individuals adding up in time. We therefore fitted the two functions to the cumulative number of new giraffe *C*_*t*_ in a Bayesian model to produce another estimate of population size (*N*_*s*_). Note that this approach assumes a constant detection rate over time, space and between individuals, given by *p*_*s*_ = *C*_20_/*N*_*s*_ and requires individuals to be uniquely identified.

#### N-mixture models

The third population size estimator we applied was the *N*-mixture model (Royle 2004). The *N*-mixture model assumes that repeated counts of animals in time and space are the outcome of combined probability models for the unknown population abundance (*N*_*N*_) and for the detection (*p*_*N*_). For population abundance, the Poisson, negative binomial and zero-inflated Poisson distributions are the most commonly used, but other discrete distributions may be considered (see below). For the detection process, a binomial distribution (with parameters *N*_*N*_ and *p*_*N*_) accounts for undetected animals. The *N*-mixture model assumes a demographically closed population and an equal detection probability for all individuals. We estimated population size by fitting four *N*-mixture models to the giraffe data (*t* = 20 days, *s* = 12 waterholes), allowing for temporal variation in detection probabilities (Kéry et al. 2009).

We replicated the analyses of population size estimation for two data sets. The first data set consisted in the number of different and uniquely recognized giraffe seen per day at each of the 12 waterholes. We used a binomial distribution to model the observation process. Here, we considered another distribution mixture accounting for the non-independence between individuals, the β-binomial–binomial *N*-mixture models (Martin et al. 2011). We discarded the zero inflated Poisson – binomial mixture because of its poor performance in general (Veech et al. 2016). For the second data set, we used the total number of giraffe seen (without individual recognition) and was hence more closely related to counts carried out in many reserves where individuals identification is not done. Here, we used a Poisson model for the observation process because double counts were very frequent from camera-trap photographs ending up with a Poisson–Poisson distribution mixture (Kéry & Royle 2020). To achieve convergence and facilitate parameter estimations, we included a temporal correlation for detection rates (first order autoregressive model, see Kéry & Royle 2020, p. 305–306). Note that in the case of Poisson – Poisson *N*-mixture models, we no longer estimate a detection probability (0 < *p* < 1) but a detection rate instead (*ψ* > 0).

We fitted all models using JAGS 4.0 (Plummer et al. 2003). We used non-informative prior distributions for all estimated parameters except for *N*_*s*_ in the rarefaction curves models, for which we used a half-normal distribution to ensure that number of animals was > 0. We ran three Monte-Carlo (MCMC) chains, with a burn-in of 10 000 iterations before saving 5 000 iterations to get the posterior distributions of parameters at convergence. We checked convergence graphically to ensure good mixing of MCMC chains and used Gelman’s *h* for an objective convergence criterion (convergence is reached when *h* is close to 1 Gelman & Pardoe 2006). The R and JAGS code we used is freely accessible on-line at https://github.com/cbonenfant.

## Results

### Camera trap data set

Giraffe were recorded at 10 of the 12 waterholes surveyed. A total of 101 individuals were identified from the camera trap images: 58 adult females, 41 adult males and two juvenile of unknown sex. For all but six individuals, we obtained identification images from both sides of the animal. For five individuals, we only had picture from the left side and only a front shot for the remaining animal. The majority of individuals (66%, *n* = 58) were seen at a single waterhole, while 27% (*n* = 24) and 7% (*n* = 6) were seen at two and three waterholes respectively. On average, 28 unique giraffes were detected per day with camera traps, with a minimum of 8 and a maximum of 54 (median of 29.5 individuals). For 98% of the individuals, we could assign the age-class.

### Population size estimates

#### Capture-recapture models

From capture histories, the best model describing the observed variability in detection rate included time variation (*i*.*e*. differences in detection probability between days), sex and whether the individual was seen at any waterhole the day before. We detected a marked variability in daily detection probabilities over the course the of the study, ranging from 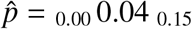 on day 2, to 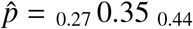 on day 15. On any day, females were 1.46 times more likely to be detected at any waterhole than males (sex effect: 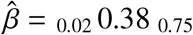). In addition, an individual was 0.63 times less likely to be detected if it was seen the day before 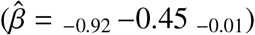. Once time, sex and previous visit had been accounted for, the remaining individual heterogeneity in detection rate was 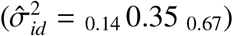. The population size estimate returned from our best model of detection rate was 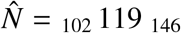 individuals (Table 1).

**Table 1.** Estimated population size 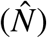 of giraffe (*Giraffa camelopardalis angolensis*) at Ongava Game Reserve, Namibia, in September 2016, from the monitoring of 10 waterholes for 21 days. The capture-recapture estimator modelled detection probability of animals accounting for daily variation, sex of individual, and whether the giraffe has previously visited a waterhole the day before or not. For the sake of comparisons, we present the detection probabilities *a posteriori* as the number of counted animals divided by 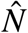. For *N*-mixture models, abundance estimation used the number of uniquely identified giraffe each day at every waterhole, hence removing double counts (Poisson–binomial and *β*-binomial–binomial mixtures) to return population size and detection probability. Another *N*-mixture model used the raw number of giraffe counted at each waterhole instead (Poisson–Poisson *N*-mixture), which is the most common configuration in wildlife counts in Africa. In this case, the model accounts for multiple counts of the same giraffe. We report here the point estimate and associated 95% credible intervals as: _lower limit_ mean _upper limit_. *C*_*v*_ stands for the coefficient of variation of 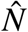.

| Estimator | $\hat{N}$ | | | $\bar{p}$ | | | $C_v$ |
| --- | --- | --- | --- | --- | --- | --- | --- |
| Capture-recapture $p_{t+h}$ | 101 | 111 | 128 | 0.79 | 0.90 | 0.99 | 6.5% |
| Capture-recapture $p_{t+sex+h}$ | 101 | 110 | 128 | 0.75 | 0.92 | 1.00 | 7.4% |
| Capture-recapture $p_{t+sex+b+h}$ | 102 | 119 | 146 | 0.69 | 0.85 | 0.99 | 6.5% |
| Rarefaction curve (Kohn) | 145 | 175 | 215 | 0.47 | 0.57 | 0.70 | 10.0% |
| Rarefaction curve (Eggert) | 104 | 117 | 134 | 0.75 | 0.86 | 0.97 | 6.3% |
| $N$-mixture (Poisson–binomial) | 173 | 215 | 263 | 0.38 | 0.47 | 0.58 | 4.4% |
| $N$-mixture (ZIP–binomial) | 173 | 215 | 263 | 0.38 | 0.47 | 0.58 | 4.4% |
| $N$-mixture (β-binomial–binomial) | 107 | 124 | 156 | 0.65 | 0.81 | 0.94 | 10.1% |
| $N$-mixture (Poisson–Poisson) | 79 | 87 | 99 | 1.02 | 1.16 | 1.28 | 5.4% |

#### Rarefaction curve models

We calculated the cumulative number of newly detected individuals over the 20 days duration of the study (Fig.2). The number of new individuals increased steeply up from day 1 to day 16 when it started to level off. It took 19 days to observe all the individuals identified during the study period (Fig.2). At the end of monitoring, 101 uniquely identified animals had been detected. Fitting the hyperbolic and exponential rarefaction curves to estimate population size gave contrasting results (Table 1). While the exponential equation returned a population size of _104_ 117 _134_ giraffe, the hyperbolic equation projected a population size 49% larger (_145_ 175 _215_). Breakage coefficients were 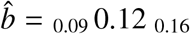 and *ĉ* = _7.9_ 11.9 _17.7_ for the exponential and hyperbolic equations respectively. Overall, the goodness-of-fit of the two rarefaction curves to the data was poor with 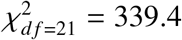 and 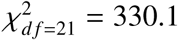 for the exponential and hyperbolic equations (Fig. 2). Precision of the estimates was of the same magnitude, close to 10% for both models (Table 1).

#### N-mixture models

We applied three different *N*-mixture models yielding contrasting results. The Poisson–binomial model returned an estimate of 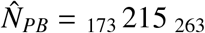 giraffes (Table 1), hence 80% larger than the estimation from the best CR model. The *β*-binomial – binomial mixture estimated abundance to 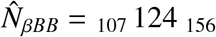 giraffes. The associated parameters of the β-binomial function were 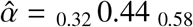 and 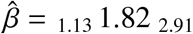, giving a correlation 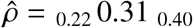. According to this model, the mean daily detection probability was 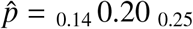, ranging between 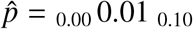 and 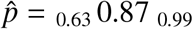. Fitting a Poisson–Poisson *N*-mixture model to raw observations led to an estimated population size of 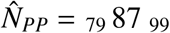 giraffe (Table 1). The mean detection rate was 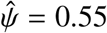 but varied from 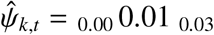 to 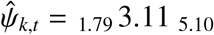 according to time and space. The first order temporal auto-correlation coefficient (AR(1)) was estimated as 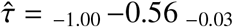. Note that the Poisson-Poisson model was particularly difficult to fit to the data as we experienced many convergence issues.

### Frequency of waterhole visits

We computed the mean time of return to a waterhole and frequency of visits from the daily probabilities as estimated from the CR model. To do so, we simulated 5 000 capture histories from a multinomial distribution which parameters were the observed detection probabilities for each day. For each capture history, we calculated the difference in days between successive visits to any waterhole, and its inverse to get the frequency of visits. The mean time of return to a waterhole of giraffe was 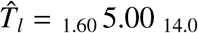 days for males, yielding a frequency of 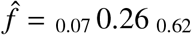. Females tended to visit waterholes more frequently with a mean time lag of 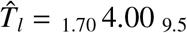 days between two observations, and a frequency 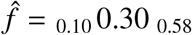 over 20 days of monitoring.

## Discussion

Population abundance is the core state variable of population dynamics from which the population growth rates is usually derived (Caughley 1977). Our study system at OGR offers a unique opportunity to apply and compare different methods to estimating giraffe abundance. Because giraffe can be recognized from their coat patterns, we were able to apply methods based on the re-observations of individuals (capture-recapture *sensu largo*), which were then compared to other abundance estimators traditionally used in wildlife monitoring in African national parks (Jachmann 2012). In comparison, the other abundance estimators deviated from −27 to +80% from the CR estimate. We caution against over-estimating giraffe abundance when using *N*-mixture models or rarefaction curves at large scale and for conservation purposes. As there is marked heterogeneity in detection probability in time and among individuals, the drinking behaviour of giraffe likely accounts for the discrepancies we report among abundance estimators, and should be carefully considered for other species monitored at waterholes. Similarly, simple counts of giraffe (i.e. without identifying individuals) at waterholes by human observers underestimate the number of giraffes by ca. −35% compared to the CR abundance estimator (77 giraffes were counted on the same period), as we might expect when detection of animals is imperfect (Seber 1982; MacKenzie et al. 2005).

Constrains, local habits and a plethora of available methods to estimate population abundance (Seber 1982) has led to inconsistent ways of monitoring wild populations of large herbivores among and, sometimes, within sites. For instance, in Hwange National Park, Zimbabwe, giraffe density estimation was derived from distance sampling (Valeix et al. 2008), while in the Serengeti, Kenya, aerial counts were preferred as an index of abundance (Strauss 2014, see also Table 1 for an overview). We show here that the choice of a particular method to estimate giraffe abundance has profound consequences on the results. On OGR, the range of estimated population sizes varied by more than two-fold, from 87 to 215, yielding densities of 0.29 and 0.71 individuals.km^−2^. Which estimator to implement and to apply to empirical data is not trivial, and comparisons of results with well-known, reference methods is advised (*e*.*g*. Corlatti et al. 2017; Pellerin et al. 2017). In our case, and in the absence of knowledge of the true number of giraffes, we considered the population size of 119 giraffe (density of 0.39 individuals.km^−2^) derived from CR models to be the most reliable among all estimates. CR methods are usually regarded as the gold standard because of their flexibility in dealing with detection probability and the long history of use since the publication of its principle by Petersen-Lincoln (Pollock 1976).

While population size as estimated from Eggert’s equation is close to CR models (117 vs. 119), the estimation from Kohn’s equation seems biologically unrealistic and should be disregarded (see also Frantz & Roper 2006, for similar results on simulated data). With 175 individuals, giraffe density (0.58 individuals.km^−2^) would be almost 3 times larger than previous estimates at Etosha National Park (Table 2), neighbouring OGR with similar rainfall conditions (Fig. 1). Such a high population density should trigger density-dependent processes, first manifested by a reduction in reproduction rates of females or low juvenile survival in large herbivores (Bonenfant et al. 2009). Rarefaction curves were shown to give biased estimation of biodiversity when species are not uniformly distributed in space (Collins & Simberloff 2009). Similarly, projecting the number of total individuals from rarefaction curves *(e*.*g*. Petit & Valiére 2006) is likely to be influenced by heterogeneity in detection probability among individuals. While Kohn’s equation returned a large number of giraffe compared to the CR estimate, Eggert’s equation almost matched our reference population size. However, with no replication of our observations and counts, we cannot assess the robustness of Eggert’s equation to heterogeneity in detection probability among individuals. All in all, the fit of the two rarefaction curves were poor (Fig. 2) making the inference on population size spurious at best.

**Table 2.** Reported densities of giraffe (*Giraffa camelopardalis ssp*.) populations in Africa (in number of individuals per km^2^). When abundance was estimated for several years, repeated lines in the same location give the range of densities recorded on the site.

| Site | Country | Density | Reference |
| --- | --- | --- | --- |
| Chobe National Park | Botswana | 0.110 (—) | Mcqualter (2018) |
| Great Rift Valley | Kenya | 0.468 (88/188) | Muller (2019) |
| Great Rift Valley | Kenya | 0.405 (77/190) | Muller (2019) |
| Mara Region | Kenya | 0.750 (—) | Ogutu et al. (2011) |
| Mara Region | Kenya | 0.080 (—) | Ogutu et al. (2011) |
| Etosha National Park | Namibia | 0.150 (—) | Brand (2007) |
| Etosha National Park | Namibia | 0.200 (—) | Brand (2007) |
| Ongava | Namibia | 0.336 (—) | This study |
| Kouré and Fandou Plateaus | Niger | 0.241 (—) | Suraud et al. (2012) |
| Lake Manyara National Park | Tanzania | 0.570 (0.570-0.580) | Kiffner et al. (2020) |
| Lake Manyara National Park | Tanzania | 1.210 (1.180-1.25) | Kiffner et al. (2020) |
| Mkomazi National Park | Tanzania | 1.165 (0.808) | Mseja et al. (2020) |
| Tarangire Ecosystem | Tanzania | 0.791 (0.073) | Lee & Bond (2016) |
| Tarangire Ecosystem | Tanzania | 1.202 (0.760) | Lee & Bond (2016) |
| Tarangire Ecosystem | Tanzania | 0.173 (0.057) | Lee & Bond (2016) |
| Saadani National Park | Tanzania | 0.100–1.400 | Treydte et al. (2005) |
| Serengeti | Tanzania | NA | Strauss (2014) |
| Shamwari | South Africa | 0.744 (—) | Hayward et al. (2007) |
| Lupande | Zambia | 1.274 (930/730) | Jachmann (2002) |
| Hwange National Park | Zimbabwe | 0.170 (—) | Valeix et al. (2008) |
| Gonarezhou National Park | Zimbabwe | 0.470 (0.140) | Ndiweni et al. (2015) |
| Malipati Sarafi Area | Zimbabwe | 0.010 (0.030) | Ndiweni et al. (2015) |
| Hluhluwe iMfolozi National Park | South Africa | 0.137 (—) | Hervé Fritz |
| Amboseli National Park | Kenya | 0.133 (—) | Hervé Fritz |
| Katavi National Park | Tanzania | 0.938 (—) | Hervé Fritz |
| Kidepo Valley National Park | Uganda | 0.037 (—) | Hervé Fritz |
| Kruger National Park | South Africa | 0.157 (—) | Hervé Fritz |
| Kruger National Park | South Africa | 0.270 (—) | Hervé Fritz |
| Mkomazi Game Reserve | Tanzania | 0.077 (—) | Hervé Fritz |
| Nairobi National Park | Kenya | 0.623 (—) | Hervé Fritz |
| Okavango Delta | Botswana | 0.250 (—) | Hervé Fritz |
| Pilanesburg National Park | South Africa | 0.286 (—) | Hervé Fritz |
| Savuti area of Chobe National Park | Botswana | 0.720 (—) | Hervé Fritz |

Although N-mixture models are more and more used to analyse count data, their reliability is regularly questioned (Dennis et al. 2015; Link et al. 2018; Knape et al. 2018; Nakashima 2020). Comparisons with other proven methods such as CR are scarce, despite their value. For giraffe, the estimation of population abundance from *N*-mixture models suffers from either a severe overestimation (215 for the Poisson–binomial) to an underestimation (87 for the Poisson–Poisson) when applied on raw, unprocessed data without identification of individuals. If individual identification is not possible, double counts are likely to occur in the raw counts. Double counting therefore be a commonly encountered situation in count operations at waterholes in many African parks. A Poisson-Poisson *N*-mixture is the natural solution to this situation by estimating a detection rate (*ψ* > 1) where individuals can be seen more than once. Unfortunately, our results suggest poor performance of the Poisson–Poisson *N*-mixture model in estimating giraffe abundance. This model produced the lowest population size estimate, being −36% smaller than CR estimate (87 vs 119 giraffe). Despite the occurrence of frequent double counts (empirical rate: 568/119 = 4.77 from CR data), the Poisson–Poisson *N*-mixture model failed to estimate this quantity correctly 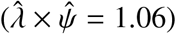, maybe because of unmodelled heterogeneity, in addition to temporal and spatial variation in the detection probability of animals. Since most giraffe live in groups, we also faced non-independence of individual detection which, when accounted for with a β–binomial distribution in the *N*-mixture model (Martin et al. 2011), returns much more sensible estimates of population size (124 individuals) than any other assumed distributions of the detection process (Table 1).

A strength of CR estimators over the rarefaction curves and *N*-mixture models is their ability to model detection probabilities not only in time and space, but also at the individual level. An important source of heterogeneity in detection probability we observed was the frequency of visit to waterholes. Giraffe visit to waterholes is primarily motivated by thirst, and if they must drink on a regular basis, they can skip drinking for several days in a row (Shorrocks 2016). On OGR, giraffe’s return frequency to waterholes was between 0.26 and 0.30 for males and females respectively (one visit every 4–5 days on average), which is lower than previously observed. For instance Shorrocks (2016) reported a frequency of 0.61, while (Caister et al. 2003) recorded daily drinking in Niger (*f* ≈ 1). Such a marked difference in drinking frequency may have both biological and technical explanations. On OGR, giraffes may find enough water in forage or access to small, non-monitored water sources, making the need to visit larger but dangerous waterholes less stringent. An alternative would be that camera traps might fail to trigger in the presence of an animal, which is sensitive to camera placement, settings and performance (Rovero et al. 2013; McIntyre et al. 2020), or because the photograph was of too low quality to allow for individual identification (*e*.*g*. blurry or dark images). Independently of its cause, this behaviour generates a particular detection pattern. Once an animal has visited a waterhole to drink, it will be less likely to be detected the following days, therefore breaking the assumption of constant detectability of many abundance estimators. In CR terminology, giraffe are “trap shy” and several solutions have been proposed by statisticians to reduce bias on abundance estimates in the CR framework (Pollock 1980).

Our study on OGR is a clear illustration that the assumption of a constant detection rate is not met, even with a fixed sampling design and a fine, daily, temporal resolution of the monitoring. Detection probability varied substantially from one day to another, ranging from 0.04 to 0.35. This result is a major warning against the use of raw (*i*.*e*. unidentified individuals) count data, such as the number of giraffes seen per day, to monitor giraffe populations in the wild (see Anderson 2001, for a general argument). Variation in daily detection probability resulted not only from the drinking and grouping behaviour of giraffe, but also from the number of camera traps in service over the course of the study. Several cameras stopped recording pictures because of battery failure or full memory cards. A sampling design based on fixed camera traps at waterholes hence does not guarantee a constant detectability. This marked variability in detection probability in time likely accounts for the discrepancy we report among the five population abundance estimators. In practice, estimating abundance of giraffe should preferably consider methods flexible enough to account for their drinking behaviour. Clearly, this may apply to other herbivore species such as zebra, greater (*T. strepsiceros*) and lesser kudu (*T. imberbis*), wildebeest (*Connochaetes taurinus*) or bushbuck (*Tragelaphus scriptus*) that all bear idiosyncratic marks.

Sampling large mammal populations with camera traps is of great practical advantage. When it comes to estimation of population abundance from camera-trap data, the long-standing issues of detection and the modelling of its heterogeneity in time, space and among individuals still apply. We found the deviation of *N*-mixture and rarefaction curve models from our reference CR estimation deteriorated when the data are not processed using individual identification. With the automation of animal re-identification using coat and fur patterns with machine learning and artificial intelligence (Miele et al. 2021), this previously time consuming processing of images data is now performed in a couple of hours. Animal identification from fur patterns is now robust, efficient, and no longer an issue for wildlife managers, and should become standard practice. We believe the gain in precision in population abundance estimation is worth the time allocated to it and will serve the conservation of many species.

## Acknowledgements

We are grateful to Jean-Michel Gaillard and Agathe Chassagneux for commenting on a previous draft of the ms and improving Fig. 1.

## Author contributions

Analysis, writing: CB, SP; study design: KS; field work: SP, KS; revision, editing: CB, SP, KS.

## Ethical standards

Of course.

